# Extraordinary Fast-Twitch Fiber Abundance in Elite Weightlifters

**DOI:** 10.1101/468744

**Authors:** Nathan Serrano, Lauren M. Colenso-Semple, Kara K. Lazauskus, Jeremy W. Siu, James R. Bagley, Robert G. Lockie, Pablo B. Costa, Andrew J. Galpin

## Abstract

Human skeletal muscle fibers exist across a continuum of slow → fast-twitch. The amount of each fiber *type* (FT) influences muscle performance but remains largely unexplored in elite athletes, particularly from strength/power sports. To address this nescience, *vastus lateralis* (*VL*) biopsies were performed on World/Olympic (female, n=6, “WCF”) and National-caliber (female, n=9, “NCF”; and male, n=6, “NCM”) American weightlifters. Participant accolades included 3 Olympic Games, 19 World Championships, 25 National records, and >170 National/International medals. Samples were analyzed for myosin heavy chain (MHC) content via SDS-PAGE using two distinct techniques: single fiber (SF) distribution (%) and homogenate (HG) composition. These athletes displayed the highest MHC IIa concentrations ever reported in healthy *VL* (23±9% I, 5±3% I/IIa, 67±13% IIa, and 6±10% IIa/IIx), with WCF expressing a notable 71±17% (NCF=67±8%, NCM=63±16%). The heavyweights accounted for 91% of the MHC IIa/IIx fibers. When compared to SF, HG overestimated MHC I (23±9 vs. 31±9%) and IIx (0±0 vs. 3±6%) by misclassifying I/IIa fibers as I and IIa/IIx fibers as IIx. These findings suggest athlete caliber (World vs. National), training experience, and body mass determine FT% more than sex and refutes the common pronouncement that women possess more slow and fewer fast-twitch fibers than men. Our results also show the abundance of pure MHC IIa and rarity of IIx in elite strength/power-trained athletes, indicate a potential link between MHC IIa/IIx frequency and body mass, and question the fidelity of HG as a measure of FT% distribution. The extreme fast-twitch abundance partially explains how elite weightlifters generate high forces in rapid time-frames. These data highlight the need for more cellular and molecular muscle research on elite anaerobic athletes.

## INTRODUCTION

Legendary Italian physician Stefano Lorenzini made the first distinction of “red” and “white” muscle fibers (myofibers) in 1678, and almost 200 years later (1873) French histologist Louis-Antoine Ranvier confirmed the existence of two distinct myofiber *types* in vertebrate skeletal muscle. Reintroduction of the skeletal muscle biopsy procedure in 1962 (1) allowed scientists to begin exploring the topic in athletes and resulted in the discovery that each fiber type (FT) is comprised of a unique myosin heavy chain (MHC) isoform signature. Human skeletal muscle therefore contains three *pure* (MHC I, IIa, and IIx) and several *hybrid* (single myofibers that co-expresses multiple MHC isoforms) FT (2). The pure and hybrid FT combine to form a robust slow → fast continuum (MHC I I/IIa IIa IIa/IIx IIx) with each displaying specific morphological, metabolic, and contractile properties (3–6). FT distribution (FT%), or the relative quantity of each FT in a given muscle, influences whole muscle function (7) and is often highly correlated with athletic performance (3, 7–13).

Extensive evidence indicates endurance athletes possess a slow-twitch myofiber majority (9, 10, 12, 14, 15), yet relatively few investigations have explored FT in speed, power, or strength athletes. Initial research in the 1970-80’s found resistance-trained men expressed high quantities (~60-65%) of fast-twitch fibers (11, 12, 15, 16), which was substantiated by later studies on elite powerlifters (17) and national-caliber (*Olympic*) weightlifters (8). This trailblazing work provided an important foundation, but used sub-elite participants (18) and/or laboratory methods that failed to accurately resolve the highly prevalent hybrids (19–21) - which compromises measurement fidelity and produces erroneous FT% conclusions (19, 22–25). More precise techniques were developed in the early 1990’s that allowed proper quantification of FT% by analyzing each single myofiber (SF).

Since this time only 13 studies (Table 1) implemented SF in young speed, power, or strength-trained individuals (5, 13, 19, 20, 22, 23, 25–30), and only 3 included females (n = 13, total). Only 5/13 included athletes: unknown-caliber male sprinters (n = 6) (25), male soccer players (n = 8) (24), elite female track and field runners (n = 6) (20), National-caliber male bodybuilders (n = 8) (19), and a former World-champion male sprinter (n = 1) (13). Accurately accounting for the full FT spectrum resulted in all five studies finding far lower MHC IIa concentrations than expected (52%, 30%, 16%, 39%, and 34%, respectively). The extremely low 16% found by Parcell et al. (2003) (20) is possibly explained by sex as females are often purported to possess more slow-twitch fibers than men (31, 32). Such sex-specific phenotypes are often the case in murine models (31), but the topic remains unexplored in athletes. Moreover, these data are difficult to interpret as the athletes sampled were from a combination of several dissimilar events (i.e., pole vault, heptathlon, 400 m hurdles, etc.).

**Table 1:** Summary of literature reporting single muscle fiber myosin heavy chain (MHC) fiber-type from the *vastus lateralis* in young speed, power, or strength-trained individuals.

| Reference | Subjects | Condition | MHC Distribution (%) |  |  |  |  |  |
| --- | --- | --- | --- | --- | --- | --- | --- | --- |
|  |  |  | I | I/IIa | IIa | IIa/IIx | IIx | I/IIa/IIx |
| Andersen (1994) | Sprinters; 6M (23y) | Post 12-week RE & Interval Training | 41 | 1 | 52 | 5 | 0 | 0 |
| Andersen (1994) | Soccer; 8M (23y) | National Players Post 12-week RE Training | 59 | 3 | 30 | 9 | 0 | <1 |
| Williamson (2001) | Non Ath; 6M (25y) 6F (21 y) | Post 12-week RE Training | 30 | 5 | 59 | 5 | 0 | 0 |
|  |  |  | 35 | 3 | 52 | 12 | 0 | 0 |
| Parcell (2003) | Track & Field; 6F (23y) | Division I / International - Caliber | 57 | 9 | 16 | 14 | 1 | 1 |
| Raue<br>(2005) | Non Ath;<br>6M (24y)<br>6M (24y) | Post-Con RE<br>Post-Ecc RE | 38<br>25 | 1<br>7 | 34<br>39 | 27<br>25 | 0<br>2 | <1<br><1 |
| Parcell<br>(2005) | Non Ath;<br>10M (22y) | Post 8-week<br>Sprint Cycle<br>Training | 34 | 8 | 44 | 12 | 0 | 2 |
| Malisoux<br>(2006) | Non Ath;<br>8M (23) | Post 8-week<br>Plyometric<br>Training | 28 | 5 | 42 | 26 | 2 | 0 |
| Kesidis<br>(2008) | Bodybuilding;<br>8M (26 y) | National-Caliber | 35 | 19 | 39 | 7 | 0 | 0 |
| Trappe<br>(2015) | Sprinter;<br>1M (? y) | Previously<br>World<br>Champion | 24 | 5 | 34 | 9 | 24 | 0 |
| Murach<br>(2016) | Non Ath;<br>9M (25y) | Resistance<br>Trained | 17 | 10 | 60 | 11 | <1 | <1 |
| Bagley<br>(2017) | Non Ath;<br>15M (25y) | Resistance<br>Trained | 20 | 10 | 58 | 11 | 1 | 1 |
| Arevalo<br>(2017) | Non Ath;<br>13M (24y) | Resistance<br>Trained | 28 | 9 | 60 | 3 | <1 | <1 |
| Tobias<br>(2017) | Non Ath;<br>1F (32y) | Concurrently<br>Trained | 45 | 13 | 31 | 9 | 0 | 2 |

MHC = Myosin heavy chain; M = Male; F = Female; y = Year; RE = Resistance exercise; Non Ath = Not a competitive athlete; Con = Concentric, Ecc = Eccentric

Numerous other knowledge gaps persist because in over 50 years of human muscle FT research only two studies have utilized SF with elite (i.e., world or international) athletes (one male sprinter and six female track and field) and no research has done so with any strength or power athlete. Thus, the purpose of this study was to examine the FT% of elite weightlifters to provide novel insight into the phenotype of competitive female and male strength and power athletes.

## METHODS

### Experimental Approach to the Problem

Twenty-one elite (*‘Olympic’*) Weightlifters (15 female, 6 male) underwent resting *vastus lateralis* biopsies between 2-96 hours after competing in either the International Weightlifting Federation World Championships or the USA Weightlifting American Open Finals (2017). All procedures and risks were explained to the athletes prior to obtaining written consent and completing medical and exercise history questionnaires. Performance records in the Snatch and Clean and Jerk (1RM), competition medals, and other accolades were gathered from personal interviews and publically available records from these or other sanctioned meets. Each muscle sample was analyzed for MHC content using two distinct FT techniques: single fiber (SF) and homogenate (HG). The University Institutional Review Board approved all experimental procedures prior to any testing.

### Participants

Participants were subdivided into three categories:

1. Olympic or World-caliber (n = 6 female): WCF
2. National-caliber female (n = 9): NCF
3. National-caliber male (n = 6): NCM

Athletes were considered “World-caliber” if they were on the most recent Olympic or World team and competed at the most recent National event. Athletes were considered “National-caliber” if they were top 5 placers at the 2017 American Open Finals meet but had never been on a World or Olympic team. Athletes spanned multiple weight categories, had a minimum of two years of National competition experience, had competed exclusively for the United States of America, and were otherwise eligible for all American National meets (Table 2). Athlete accolades at the time of data collection included participation in 3 Olympic Games, 19 World Championships, 11 Pan American Championships, 49 National Championships, 32 American Opens, 8 University National Championships, and 25 Junior World/Pan American/National Championships. Participants also held 25 National records and >170 National/International medals. One athlete had tested positive for substances prohibited by the World Anti-Doping Agency and was suspended from the sport for two years prior to participating in the study.

**Table 2:**
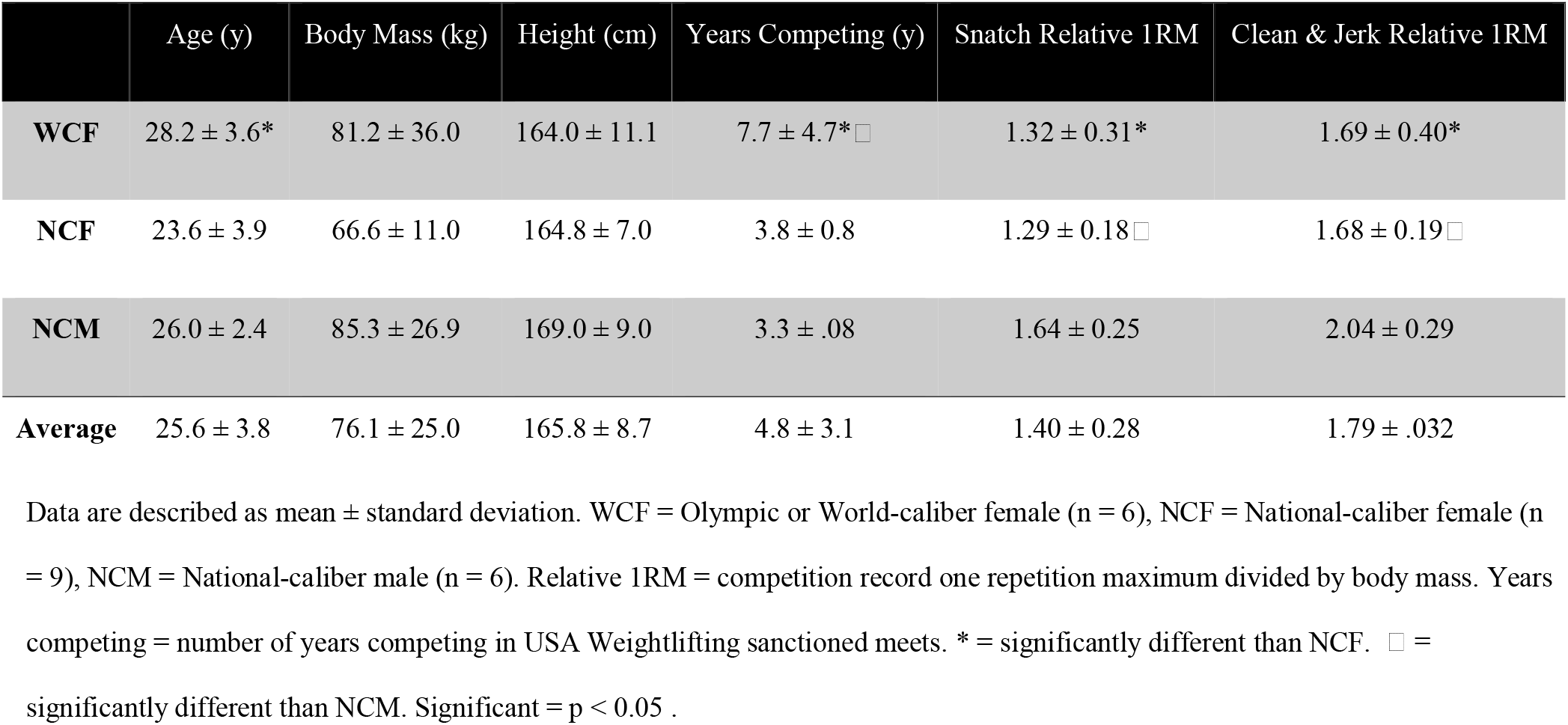
Descriptive information of elite female and male American weightlifters.

|  | Age (y) | Body Mass (kg) | Height (cm) | Years Competing (y) | Snatch Relative 1RM | Clean & Jerk Relative 1RM |
| --- | --- | --- | --- | --- | --- | --- |
| **WCF** | 28.2 ± 3.6* | 81.2 ± 36.0 | 164.0 ± 11.1 | 7.7 ± 4.7*□ | 1.32 ± 0.31* | 1.69 ± 0.40* |
| **NCF** | 23.6 ± 3.9 | 66.6 ± 11.0 | 164.8 ± 7.0 | 3.8 ± 0.8 | 1.29 ± 0.18□ | 1.68 ± 0.19□ |
| **NCM** | 26.0 ± 2.4 | 85.3 ± 26.9 | 169.0 ± 9.0 | 3.3 ± .08 | 1.64 ± 0.25 | 2.04 ± 0.29 |
| **Average** | 25.6 ± 3.8 | 76.1 ± 25.0 | 165.8 ± 8.7 | 4.8 ± 3.1 | 1.40 ± 0.28 | 1.79 ± .032 |

Data are described as mean ± standard deviation. WCF = Olympic or World-caliber female (n = 6), NCF = National-caliber female (n = 9), NCM = National-caliber male (n = 6). Relative 1RM = competition record one repetition maximum divided by body mass. Years competing = number of years competing in USA Weightlifting sanctioned meets. \* = significantly different than NCF. □ = significantly different than NCM. Significant = p < 0.05 .

### Procedures

#### Vastus Lateralis Muscle Biopsies

Following 30 minutes of supine rest, athletes underwent a mid-muscle belly (approximately halfway between the greater trochanter and patella) biopsy of the *vastus lateralis*. A detailed description of the biopsy procedure has been previously described by our lab (9, 22, 23, 33). Briefly, a small area of the thigh was numbed by injection of a local anesthetic (Xylocaine/Lidocaine without epinephrine). An approximately ¼ inch incision was made in the superficial cutaneous tissues. Muscle samples were obtained using the Bergström technique with suction (1), immediately cleansed of excess blood and connective tissue, divided into approximately 10-15 mg strips, placed into cold skinning solution (125 mM K propionate, 2.0 mM EGTA, 4.0 mM ATP, 1.0 mM MgCl_2_, 20.0 mM imidazole [pH 7.0], and 50% [vol ml/vol ml] glycerol), and stored at −20° C for at least one week. Each sample was split such that a portion (~5 mg) could be used for single fiber isolation or homogenization. The incision site was cleaned, pulled closed with a sterile Band-Aid, and covered with sterile gauze and cohesive bandage tape.

#### Myosin Heavy Chain Fiber Type Identification

All biopsy samples were analyzed for MHC via SDS-PAGE using two distinct techniques: single fiber *distribution* (SF) and homogenate *composition* (HG). For SF, individual fibers (N = 2,147; 102 ± 3 fibers per athlete) were mechanically isolated with fine tweezers under a light microscope and placed in 80 μl of sodium dodecyl sulfate (SDS) buffer (1% SDS, 23 mM EDTA, 0.008% bromophenol blue, 15% glycerol, and 715 mM b-mercaptoethanol [pH 6.8]). HG samples (~5 mg) were hand homogenized and then diluted between 1:10 to 1:50 based on sample amount and protein quantity. As described in detail elsewhere (5, 9, 22, 23, 27), 1-2 μl aliquots of both SF or HG (run separately) were then loaded into individual wells in a 3.5% loading and 5% separating gel (SDS-PAGE), run at 5°C for 15.5 hours (SE 600 Series; Hoefer, San Francisco, CA, USA), and silver stained for MHC identification. The SF approach used known molecular weights and standards to identify the MHC isoform (MHC I, I/IIa, IIa, IIa/IIx, and IIx) of each individual myofiber. This enabled the most accurate calculation of the distribution/percent frequency (FT%) of each FT contained within the muscle sample (21). For example, if 100 fibers were analyzed and 30 were identified as MHC I, 60 as MHC IIa, and 10 as MHC IIa/IIx, the FT% would be calculated as 30% MHC I, 60% MHC IIa, and 10% MHC IIa/IIx. HG utilized densitometry (ImageJ, National Institutes of Health, Bethesda, MD) to quantify the relative MHC protein composition (i.e., percent area occupied by each pure isoform; MHC I, IIa, and IIx) of each sample, which is correlated highly with FT area (34). Thus, SF indicates how frequently each isoform exists but cannot address how much area each FT occupies within the muscle. HG addresses the latter, but cannot delineate hybrids, therefore inaccurately quantifying FT% (9, 21–25, 27).

### Statistical Analysis

Potential differences between groups in descriptive information were examined via ANOVA. For SF, potential differences in FT% between groups were assessed via a 3 (group: WCF, NCF, NCM) x 4 (fiber type: MHC I, I/IIa, IIa, IIa/IIx) ANOVA. For HG, potential differences in FT composition between groups were examined via a 3 (group: WCF, NCF, NCM) x 3 (fiber type: MHC I, IIa, IIx) ANOVA. Comparison of SF vs. HG was accomplished by a 2 (group: SF, HG) x 3 (fiber type: MHC I, IIa, IIx) ANOVA. Effect size was calculated with Cohen’s D (0.2 = small difference, 0.5 = medium difference, and 0.8 = large difference) to identify the magnitude of difference between two groups. Pearson Product Moment Correlations (r) were assessed for WCF, NCF, and NCM between 1RM, body mass, and SF FT%. All individual FT data are reported in Table 3. Data are reported as mean ± standard deviation (SD), unless otherwise noted. Significance was established *a priori* at an alpha level of *p < 0.05*. All analyses were performed with SPSS (SPSS Statistics Version 24, IBM).

**Table 3.**
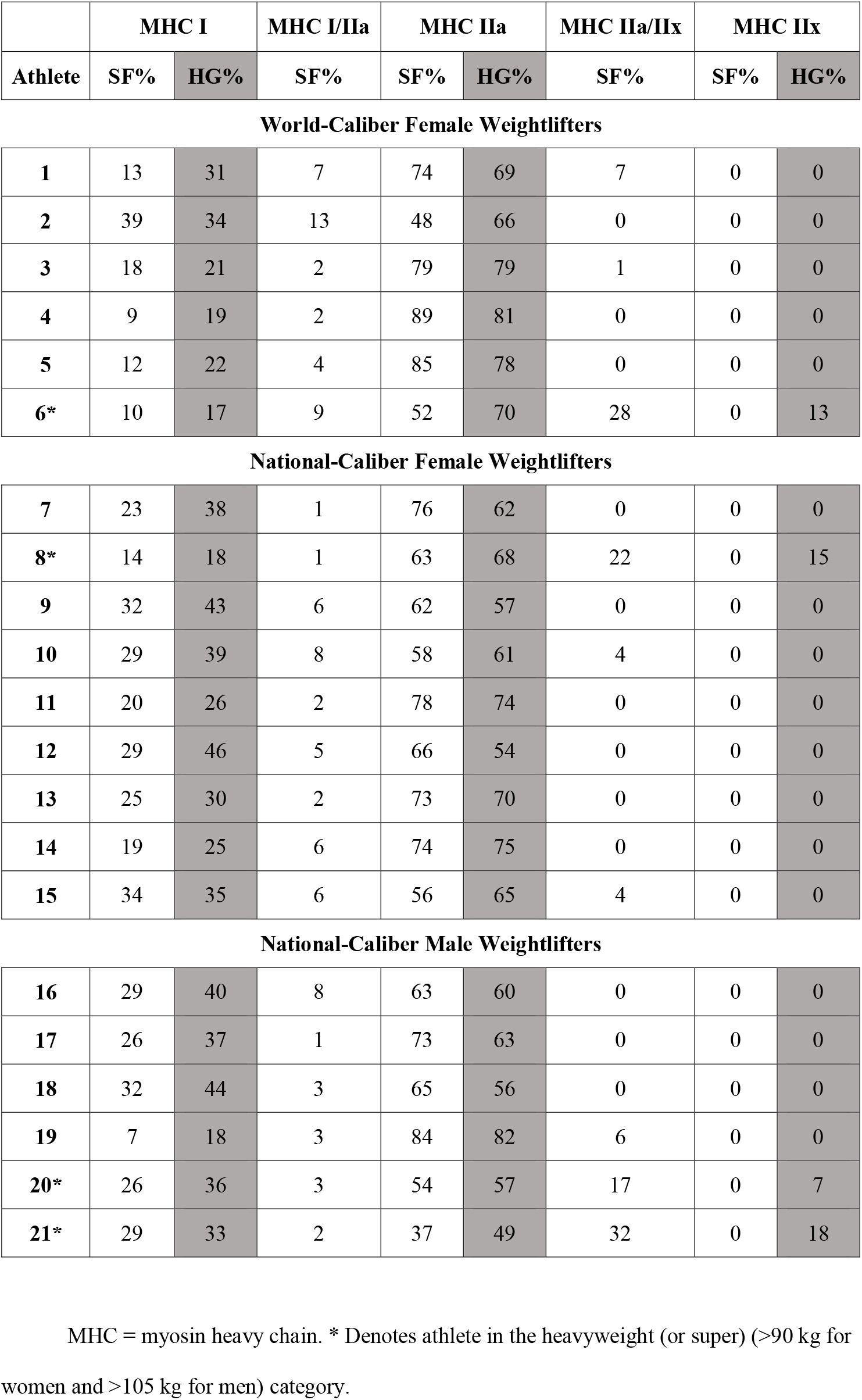
Individual single muscle fiber type distribution (SF) and homogenate composition (HG) of elite female and male American weightlifters. Data are reported as a percentage.

|  | MHC I |  | MHC I/IIa | MHC IIa |  | MHC IIa/IIx | MHC IIx |  |
| --- | --- | --- | --- | --- | --- | --- | --- | --- |
| Athlete | SF% | HG% | SF% | SF% | HG% | SF% | SF% | HG% |
| **World-Caliber Female Weightlifters** |  |  |  |  |  |  |  |  |
| **1** | 13 | 31 | 7 | 74 | 69 | 7 | 0 | 0 |
| **2** | 39 | 34 | 13 | 48 | 66 | 0 | 0 | 0 |
| **3** | 18 | 21 | 2 | 79 | 79 | 1 | 0 | 0 |
| **4** | 9 | 19 | 2 | 89 | 81 | 0 | 0 | 0 |
| **5** | 12 | 22 | 4 | 85 | 78 | 0 | 0 | 0 |
| **6\*** | 10 | 17 | 9 | 52 | 70 | 28 | 0 | 13 |
| National-Caliber Female Weightlifters |  |  |  |  |  |  |  |  |
| **7** | 23 | 38 | 1 | 76 | 62 | 0 | 0 | 0 |
| **8\*** | 14 | 18 | 1 | 63 | 68 | 22 | 0 | 15 |
| **9** | 32 | 43 | 6 | 62 | 57 | 0 | 0 | 0 |
| **10** | 29 | 39 | 8 | 58 | 61 | 4 | 0 | 0 |
| **11** | 20 | 26 | 2 | 78 | 74 | 0 | 0 | 0 |
| **12** | 29 | 46 | 5 | 66 | 54 | 0 | 0 | 0 |
| **13** | 25 | 30 | 2 | 73 | 70 | 0 | 0 | 0 |
| **14** | 19 | 25 | 6 | 74 | 75 | 0 | 0 | 0 |
| **15** | 34 | 35 | 6 | 56 | 65 | 4 | 0 | 0 |
| National-Caliber Male Weightlifters |  |  |  |  |  |  |  |  |
| **16** | 29 | 40 | 8 | 63 | 60 | 0 | 0 | 0 |
| **17** | 26 | 37 | 1 | 73 | 63 | 0 | 0 | 0 |
| **18** | 32 | 44 | 3 | 65 | 56 | 0 | 0 | 0 |
| **19** | 7 | 18 | 3 | 84 | 82 | 6 | 0 | 0 |
| **20\*** | 26 | 36 | 3 | 54 | 57 | 17 | 0 | 7 |
| **21\*** | 29 | 33 | 2 | 37 | 49 | 32 | 0 | 18 |

MHC = myosin heavy chain. \* Denotes athlete in the heavyweight (or super) (>90 kg for women and >105 kg for men) category.

## RESULTS

### Descriptive

WCF were significantly older than NCF, but not NCM (Table 2). WCF also had significantly more years of sport competition experience than NCF and NCM. Yet, NCM exceeded both WCF and NCF in relative strength in both the Snatch 1RM and Clean and Jerk 1RM.

### Single Fiber Analysis (SF)

FT% for all lifters combined was 23 ± 9% I, 5 ± 3% I/IIa, 67 ± 13% IIa, and 6 ± 10% IIa/IIx. No MHC IIx or I/IIa/IIx fibers were identified. No significant differences existed between groups, despite WCF possessing 8% (absolute, not percent difference) less MHC I than NCF (d = 0.88) and NCM (d = 0.78) (Figure 1). The difference in MHC IIa between WCF and NCM (also 8%) was also not statistically significant, but had a moderate effect size (d = 0.50). The vast majority of the MHC IIa/IIx fibers (91%) belonged to just five lifters, all of whom competed in the heavyweight or super heavyweight categories (≥ 90 kg for women and ≥ 105 kg for men). Thus, significant correlations were present between body mass and MHC IIa/IIx frequency for WCF (*r* = 0.919, p = 0.010) and NCF (*r* = 0.826, p = 0.006) while a trend existed for NCM (*r* = 0.757, p = 0.080).

**Figure 1.**
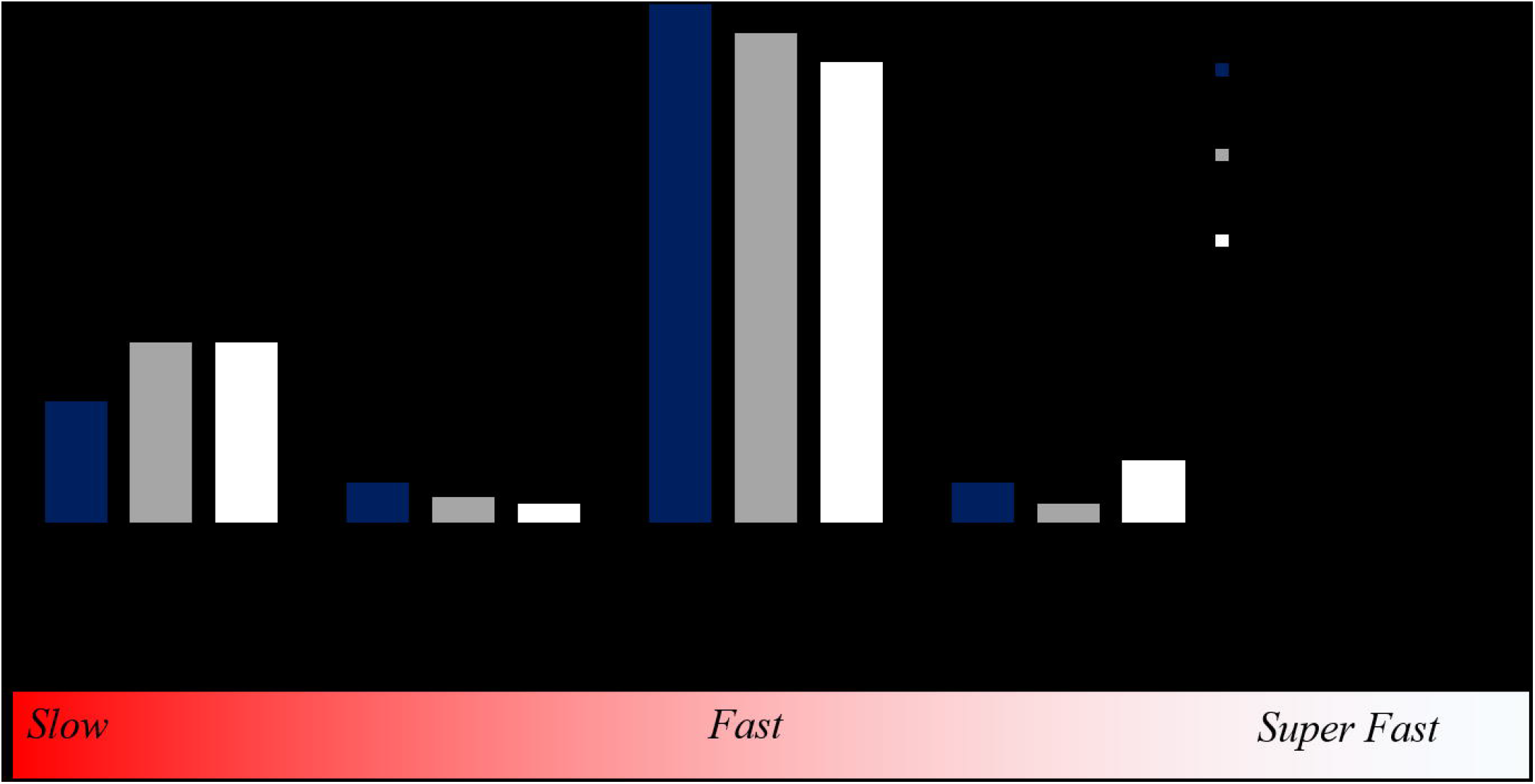
Myosin heavy chain (MHC) fiber type distribution of elite American weightlifters. WCF = Olympic or World-caliber female (n = 6), NCF = National-caliber female (n = 9), NCM = National-caliber male (n = 6). Data are reported as a percentage ± standard deviation.

### Homogenate Analysis (HG)

FT composition for all lifters combined was 31 ± 9% I, 67 ± 9% IIa, and 3 ± 6% IIx. MHC I tended (p = 0.08) to be lower in WCF (24 ± 7%) than NCF (33 ± 9%, p = 0.125, d = 1.33) and NCM (35 ± 9%, p = 0.106, d = 1.14), yet MHC IIa was significantly higher (p = 0.046) in WCF (74 ± 6%) than NCF (65 ± 7%, p = 0.145, d = 1.28) and NCM (61 ± 11%, p = 0.043, d = 1.39). FT was significantly different (p < 0.001) between SF and HG for MHC I (p = 0.005) and MHC IIx (p = 0.046), but not MHC IIa. SF MHC IIa/IIx and HG MHC IIx were highly correlated (*r* = 0.96, p < 0. 001). No correlations existed for SF or HG between FT% and Snatch or Clean and Jerk relative 1RM.

## DISCUSSION

The current study resulted in the most detailed investigation of muscle phenotype in Olympic and World-caliber anaerobic athletes published to date. Additionally, the data enabled the first comparison and differentiation of World vs. National-caliber athletes at the single fiber level. The current study was also the most precise description of FT% in strength or power sport competitors, and the first ever in females. The MHC IIa abundance was the highest in healthy muscle (VL) ever reported, especially for females. These data suggest athlete caliber and/or training history influences FT% more than sex *per se* and also questions the pronouncement that male athletes possess more fast-twitch myofibers than females. Our utilization of two different typing methods confirmed the limitations of HG for FT% (inappropriately categorizes MHC I/IIa as MHC I and MHC IIa/IIx as MHC IIx) and also allowed identification of a previously undocumented relationship between body mass and MHC IIa/IIx concentrations. The unique morphology and phenotypes in our participants highlight the need to further study elite anaerobic athletes, particularly females.

WCF contained the highest concentration of MHC IIa (71%) ever reported in the literature to our knowledge. NCF (67%) and NCM (63%) also possessed more MHC IIa than previous research in competitive bodybuilders (40%) (16, 19) as well as power/weightlifters (8, 11, 12, 15, 16, 18), elite track and field athletes (20, 35), and resistance-trained men (18, 19, 22, 23, 27, 29, 36), which all ranged from 50-60%. Only six previous studies using SF have found pure MHC IIa concentrations of >50%, with just two reporting 60% (Table 1). The resulting minimal MHC I (~17-25%) in our athletes was strikingly lower than elite female track and field athletes (57%) (20) and National-caliber bodybuilders (35%) (19). These pronounced differences are likely explained by the substantial dissimilarities in training styles (e.g., external loading strategies, contraction type and velocity, training frequency, etc.) between the various sports. More research is therefore needed to continue delineating the subtle but significant differences in FT% between top-performing athletes in various anaerobic sports and the specific role each training approach might play on altering MHC I and IIa distribution. Although it did not reach statistical significance, large ES were evident and MHC IIa frequencies of 74%-89% occurred in 66% of WCF but only in 44% and 33% of NCF and NCM, respectively. Thus, scientists should continue to examine what separates World from National-level athletes.

WCF differed from NCF and NCM in both sex and years competing in the sport (~8 vs. 3 y). Sex comparisons in athletes remains tenuous (31, 37) because nearly all investigations utilize non-gold standard FT% methods (21) and sedentary (38, 39) or “recreationally active” individuals (32). Not only do our findings contradict the claim that women possess more slow-twitch myofibers than men (40), they illustrate the opposite when accounting for talent level (WCF < NCF = NCM). The current cross-sectional study-design precludes direct analysis, but extensive research affords strong support for training history as a critical determinate of FT% (2, 9, 26, 28, 30, 41–44). Chronic exercise generally decreases hybrids (30, 42) and induces style-specific shifts in FT% such as increases in MHC I with endurance (9, 43) or MHC IIa with sprint (28), plyometric (26), or strength training (36, 43–45). For example, MHC I concentrations in an individual with extensive endurance exercise history were nearly double that of his non-exercising monozygous twin (9). Another study reported an increase in MHC IIa from 46% to 60% following 19 weeks of resistance training (36). MHC IIa/IIx fibers appear particularly responsible for exercise-induced increases in MHC IIa and are thus uncommon in exercise-trained individuals (9, 20, 22–25, 29, 43). A reduction of MHC IIx in favor of IIa following chronic resistance exercise is also purported extensively in the literature (28, 34), yet the overwhelming majority of this evidence comes from experiments with methodologies directly shown here and elsewhere (9, 24, 25) to produce erroneous FT% conclusions.

Most research from the 1970’s – 2000’s utilized either ATPase histochemistry or HG SDS-PAGE to determine FT% (8, 14–17, 24, 25, 34, 36). Similar to SF, histochemistry allows assessment of individual fibers for calculation of percent distribution, yet it does not enable simultaneous delineate of hybrids (36). HG suffers the same drawback and actually indicates FT area/composition (34) more so than distribution making it greatly influenced by the size of each fiber; which is not uniform across all FT (particularly in resistance trained individuals) (46). All three approaches hold strong merit and are often correlated to each other (34, 47) and performance (8), but are clearly not interchangeable for maximally precise FT% assessment. In the current study, HG accurately quantified MHC IIa (within 0-4%), but not I or IIx. MHC I was overestimated by 8% percent (23 vs. 31%), which is largely explained by the non-differentiated MHC I/IIa fibers (5%). HG also greatly exaggerated MHC IIx, particularly in individuals with >4% MHC IIa/IIx. The inability of HG to account for MHC IIa/IIx explains why MHC IIx appear common in some studies (48) even though their actual abundance in healthy human skeletal muscle is extraordinarily rare; typically <0.1% (9, 22–25, 27) and 0 of the >2,100 isolated fibers from the current sample. Thus, the seeming conversion of MHC IIx to IIa with exercise is more precisely IIa/IIx changing to IIa.

For these reasons MHC IIa/IIx are typically inversely associated with muscle health and physical activity (9, 43). Yet, the heavyweights (male and female) expressed irregularly high concentrations (24%) and accounted for 91% of all MHC IIa/IIx myofibers. Terzis and colleagues (2010) noted a similar abnormal abundance of MHC IIx (typed via HG, so likely IIa/IIx) in six large (116 kg, body fat composition >22%), but presumably well strength-trained throwers (35). Body composition was not assessed in the current study and little research exists on well-trained, but obese individuals. Thus, additional studies across a broader spectrum of physical size are required to truly interpret the correlations between body mass and MHC IIa/IIx prevalence.

Another juxtaposition was that of FT% and performance. Previous work in 94 kg male competitive weightlifters found strong correlations between FT composition (HG) and percent FT area to both snatch 1RM and vertical jump height (8), but not clean and jerk 1RM. We failed to identify any such correlations, but also utilized multiple sexes and weight classes. Thus, while FT% differed between our groups, that factor alone did not predict performance among our lifters. Several possible explanations exist for this discrepancy. First, FT area may determine whole muscle strength more than FT%. Second, neither studies found correlations to the clean and jerk, which is heavier and slower than the snatch or vertical jump. This compliments previous isokinetic research (23) and indicates FT% does not predict performance on strength tasks among strength-trained individuals. FT% probably determines movement speed more than force production (7). Further speculation on this point is unwarranted as limitations prohibited the ability to assess FT-specific size or contractile properties, which likely differed significantly across our groups (49) and are known to changes with training (3, 46).

## CONCLUSION

This study provides novel insight into the muscle phenotype of elite competitive strength and power athletes. Our data indicate athlete caliber, training history, and body mass dictate FT% more than sex *per se*, but more work is needed to draw firm conclusions. The extreme fast-twitch abundance partially explains how elite weightlifters are able to generate high forces in short time-frames. Most athletes contained few hybrids and no MHC IIx or I/IIa/IIx, except the heavyweights who possessed large quantities of IIa/IIx. Future research should use high fidelity techniques to explore FT-specific distribution, size, and contractile properties in female and male athletes of various caliber, sports, and body size; ideally across several years of competition. The resulting knowledge could have practical significance if it enabled experimentation of differing training volumes or recovery protocols based on athlete-specific FT properties (50). More detailed fiber type profiles of elite strength and power athletes may eventually enable strength and conditioning professionals to create more individualized and effective programs.

## ACKNOWLEDGEMENTS

Funding for this project was provided by Renaissance Periodization. The authors would like to thank Irene S. Tobias and Cameron Yen for their help with this project. Contact the corresponding author at.

